# ERK5 inhibition triggers CDK6 proteasomal degradation and enhances palbociclib efficacy in cancer cells

**DOI:** 10.64898/2026.09.17.752291

**Authors:** Claudia Faundez-Vidiella, Ylenia Sfragano, Ignazia Tusa, Barbara Stecca, Sergio Espinosa-Gil, José M Lizcano, Elisabetta Rovida

**Author notes:** Corresponding authors: Elisabetta Rovida, Viale GB Morgani, 50 –50134 Firenze, Italy,; Sergio Espinosa-Gil, Vall Hebron Institut de Recerca, (VHIR), Passeig Vall d’Hebron, 119-129, 08035 Barcelona, Spain. C. Faundez-Vidiella and Y. Sfragano should be considered joint first authors. J.M. Lizcano and E. Rovida should be considered joint senior authors. (C.F-V), (J.M.L.), (S. E-G), (Y.S.), (I.T.), (E.R.), (B.S.).

## Abstract

Advanced endometrial cancer (EC) and melanoma are two malignancies with poor treatment options at advanced stages. CDK4 and CDK6 kinases play a central role in the regulation of cell proliferation by controlling progression through G1/S transition. CDK4/CDK6 inhibitors are currently in clinical trials for advanced ECs, whereas melanoma tumors frequently carry mutations affecting the CDK4/CDK6 pathways that support the therapeutic potential of these kinases. The MAP kinase ERK5 promotes tumor progression by driving cell-cycle progression, yet its functional interplay with CDK4/6-dependent cell cycle control remains poorly defined.

Here, we investigated the relationship between ERK5 and CDK4/6 in melanoma and serous EC cells, uncovering the benefits of their co-targeting. Both genetic and pharmacological inhibition of ERK5 induced CDK6 proteasomal degradation, without affecting CDK4 protein levels. Co-treatment with the ERK5 inhibitor JWG-071 and palbociclib synergistically reduced cell viability and increased apoptosis in both melanoma and EC cells, compared with single-agent treatments. Mechanistically, combined inhibition of ERK5 and CDK4/6 reinforced cell-cycle inhibitory signaling through p21 induction and reduced phospho-retinoblastoma levels. These findings suggest that targeting ERK5 may improve the anticancer efficacy of palbociclib, at least in melanoma and serous EC tumors.

## INTRODUCTION

Endometrial cancer (EC) is the fourth most common neoplasia in women^1^. Type I EC is commonly hormone-dependent and is associated with favorable outcomes^2^. For advanced type I EC, the combination of hormonal therapy and cell cycle inhibitors (such as the CDK4/6 inhibitor palbociclib) are currently in phase 2b clinical trials^3,4^. In contrast, type II ECs, including serous EC, are non-hormone dependent and have a dismal prognosis^5^, making new therapeutic strategies an urgent need. Melanoma is among the deadliest skin cancer, with a poor prognosis in the advanced stages^6^. Up to 90% of melanomas harbor genomic alterations affecting components of the CDK4/CDK6-Rb pathway^7^, suggesting a possible use of CDK4/CDK6 inhibitors for its treatment^8^.

The Extracellular signal-Regulated Kinase 5 (ERK5) is a Mitogen-Activated Protein Kinase (MAPK) that is ubiquitously expressed in mammalian tissues, and it is activated in response to growth factors, cytokines, and different forms of stress by direct phosphorylation of its only upstream activator MEK5^9^. The MEK5-ERK5 pathway is an oncogenic pathway that contributes to tumor development. Overactivation of the pathway, or increased ERK5 or MEK5 protein levels, correlates with tumor aggressiveness and poor prognosis of many solid tumors^10^, including prostate^11^, EC^12^, breast cancer^13^, melanoma^14^ and lung cancer^15,16^. Several studies have shown that genetic and pharmacological inhibition of ERK5 impairs cancer cell proliferation and viability, both as monotherapy and in combination with standard anticancer therapies^17,18^. In the context of targeted therapies, ERK5 inhibitors potentiate the anticancer activity of the BRAF-V600E inhibitor vemurafenib in human melanoma cell lines and xenograft models^14^, as well as of Smoothened or GLI inhibitors (MRT-92 and GANT61) in 2D and 3D cultures of human melanoma cells^19,20^. Moreover, ERK5 inhibitors sensitize EC cells and tumor xenografts to paclitaxel^12^, colorectal cancer cells and tumor xenografts to 5-fluorouracil^21^, and diverse cancer cell models to extrinsic apoptosis exerted by death receptor agonist and natural killer cells^22^.

ERK5 plays an important role in cell-cycle progression at the G1/S phase transition, where ERK5 promotes cyclin D1 transcription^23^. Notably, ERK5 pathway inhibition increases the levels of the cyclin-dependent kinase inhibitors such as p21 or p27^24,25^, through regulating their stability at mRNA level or protein level^26^. Consistently, ERK5 inhibitors impair retinoblastoma (Rb) phosphorylation in mouse embryonic fibroblasts and melanoma cells, leading to cell cycle arrest at the G1/S or G2/M phases in a cell type-dependent manner^26–28^. Based on these observations, we hypothesized that ERK5 inhibition could enhance the efficacy of the clinically approved CDK4/6 inhibitor palbociclib in cancer types with unmet therapeutic needs, such as serous EC and melanoma.

## MATERIALS AND METHODS

### Cell cultures

Human serous endometrial cancer cell lines ARK1 (RRID: CVCL_IV72) and ARK2 (RRID: CVCL_IV73), and human metastatic melanoma BRAFV600E cell line A375 (RRID: CVCL_0132) were purchased from ATCC (Manassas, VA, USA). Triple wild-type (i.e. lacking BRAF, N-RAS and NF1 mutations) SSM2c metastatic melanoma cells have been previously described^29^. Cells were cultured in Dulbecco’s modified Eagle’s medium (DMEM) supplemented with 10% fetal bovine serum (FBS), 2 mM glutamine, and 1% penicillin/Streptomycin (Euroclone, Paignton, UK) (complete media). Cells were maintained at 37°C in a humidified atmosphere containing 5% CO2. Cell lines were yearly authenticated (Promega PowerPlex Fusion System kit; BMR Genomics s.r.l, Padova, Italy). The presence of mycoplasma was periodically tested by PCR.

### Drugs

The ERK5 inhibitor JWG-071^30^ and the proteasome inhibitor MG-132 were from Sigma-Aldrich (St. Louis, MO)^31^. The CDK4/CDK6 inhibitor palbociclib was from MedChemExpress (Monmouth Junction, NJ, USA).

### RNA interference

Stable knockdown of ERK5 in melanoma and EC cells was performed using lentiviral vectors as previously reported^24^. Lentiviral vectors (TRC1.5-pLKO.1-puro) carrying an shRNA non-targeting sequence or targeting human MAPK7 (shERK5-1 NM_139032.X, clone ID: TRCN0000010262; shERK5-2 NM_139032.X, clone ID: TRCN0000010275) were used. Infected cells were selected with 2 µg/mL puromycin for at least 72 hours. Cells were lysed for further purposes after 5 days.

### Cell lysis and Western Blot

Cells were lysed as previously described^32^, using cold radioimmunoprecipitation assay (RIPA) buffer, (25mM Tris-HCl pH 7.9; 150mM NaCl; 1mM EGTA; 5mM sodium pyrophosphate; 0.05% w/v deoxycholic acid; 0.1% w/v SDS; 1% w/v NP-40). Cell lysates were sonicated (4x10s) and centrifuged at 12.000 rpm for 12 minutes at 4°C to remove DNA and cellular debris from the sample.

Proteins were separated by SDS–PAGE and transferred onto nitrocellulose membranes by electroblotting. Membranes were blocked and incubated with antibodies as previously described^33^. Infrared imaging (Odyssey, LI-COR Biosciences, Lincoln, NE, USA) detection was performed. Antibodies are listed in Table 1.

### MTT assay

Cell viability was quantified using 3-(4,5-dimethylthiazol-2-yl)-2,5-diphenyltetrazolium bromide (MTT; Sigma-Aldrich, St. Louis, MO). 10.000 ARK1 cells/well, 7.000 ARK2 cells/well, 4.000 A375 cells/well, and 6.000 SSM2c cells/well were seeded in 48-well plates, and after 24 hours they were treated with the indicated drugs. After 72 hours, 250 μL of complete media containing 0.5 mg/mL MTT were added to each well. Plates were incubated for 50 minutes at 37°C, and MTT solution was aspirated and 100 μL of DMSO was added to each well. Absorbance was measured in an Epoch reader (Agilent Technologies, Santa Clara, CA) using a 560 nm reading wavelength and a 620 nm reference wavelength.

### Cell counting by optical microscopy

In p60 plates, 800000 ARK1, 500000 ARK2, 300000 A375 or 300000 SSM2c cells were seeded in complete media and 24 hours later cells were treated with the different drugs for 24 hours. Images were captured using an Olympus CKX41 microscope (Leica Microsystems, Mannheim, Germany) at 10x magnification.

### Determination of apoptosis

To quantify apoptosis, cells were detached using non-enzymatic acutase solution (Euroclone, Paignton, UK) and then centrifuged, resuspended in binding buffer and incubated with FITC-labelled Annexin-V (Roche Diagnostics, Basel, Switzerland) and propidium iodide (PI) for 15 min at room temperature (RT) in the dark, according to manufacturer’s protocol. Flow cytometry was performed using a FACSCanto (Becton–Dickinson, Franklin Lakes, NJ, USA), as previously reported^34^.

### Statistical analysis

Statistical analyses were performed using the GraphPad Prism 5 software (RRID:SCR_002798) with T-student and one-way ANOVA variance analysis, followed by Tukey’s multiple comparison test. Western blot protein quantification was performed using ImageJ (RRID:SCR_003070) software. Synergy analysis was performed using BLISS algorithm on the Synergy Finder 3.0 platform. Statistical significance cut-off was established as p<0.05. Significance values are expressed as: *, p<0.05; **, p<0.01; ***, p<0.001; ****, p<0.0001.

## RESULTS

### Loss of ERK5 reduces CDK6 protein levels

To assess the role of ERK5 as a potential regulator of CDK proteins involved in G1/S phase transition, we first studied the effect of ERK5 inhibition on CDK4 and CDK6 protein levels. For this purpose, we investigated the effect of the ERK5 inhibitor JWG-071^12,30^ on two human high-risk serous EC (ARK1 and ARK2) and two human metastatic melanoma (A375 and SSM2c) cell lines. Pharmacological inhibition of ERK5 led to a significant impairment of CDK6 protein levels across all four cell lines, without affecting those of CDK4 protein (Fig. 1A-B). This effect was also observed for other ERK5 inhibitors (ERK5i) such as BAY-885 and AX15836 (Supplementary Fig. 1). Consistently, ERK5 silencing using two different specific shRNAs resulted in a significant decrease in CDK6 protein levels in both tumor models (Fig. 1C), further supporting a role of ERK5 in regulating CDK6 protein levels.

**Figure 1.**
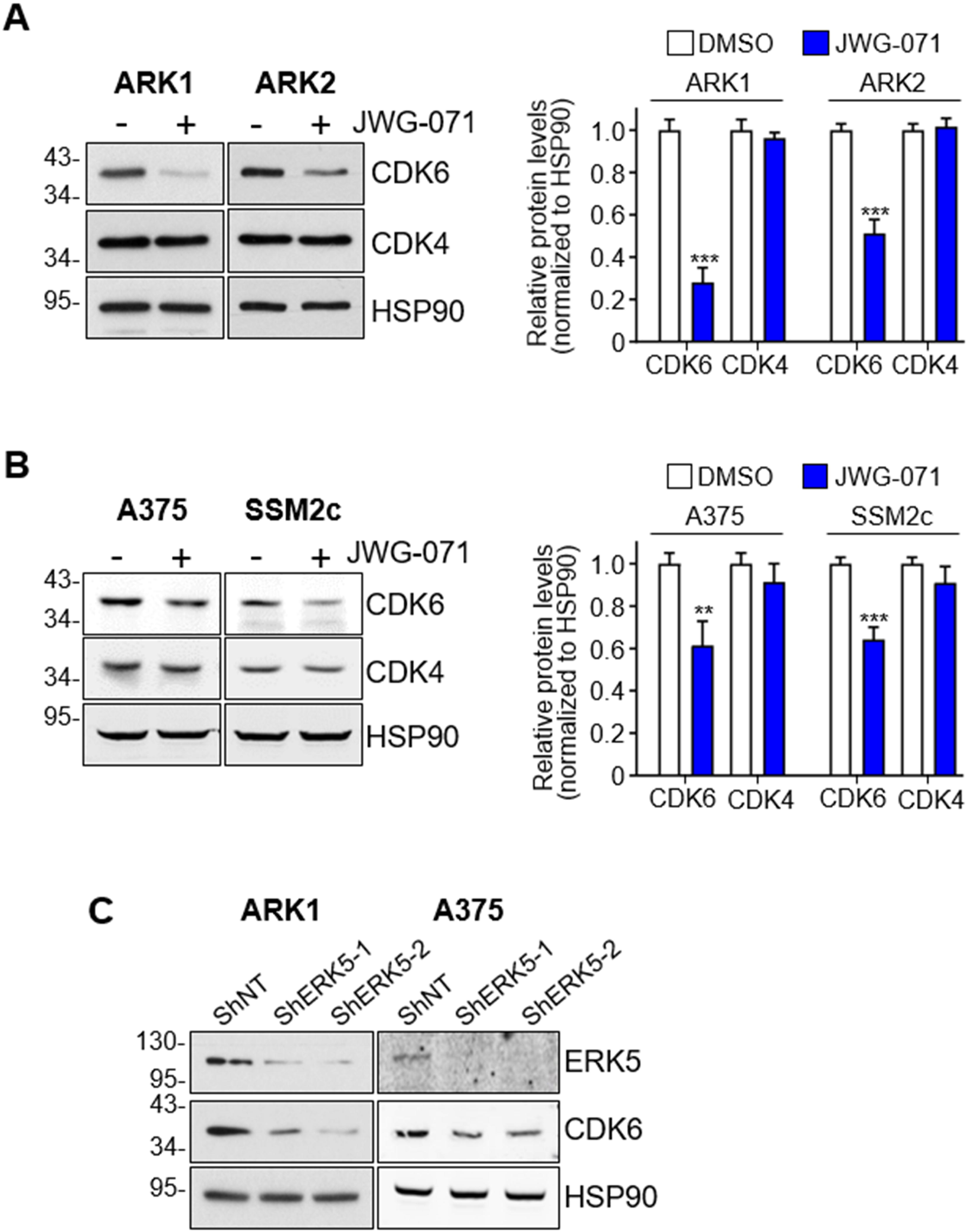
ERK5 inhibition reduces CDK6 protein levels. (**A, B**) Human serous EC ARK1 and ARK2 (A) or human melanoma A375 and SSM2c (B) cells were treated with 5μM JWG-071 for 24 hours. Cells were then lysed and Western Blot was performed with the indicated antibodies. Molecular weight markers are indicated on the left. The graphs show average densitometric values normalized on HSP90 content ± SD of three independent experiments. One-way ANOVA variance analysis was used to assess statistical significance **, p<0.01, ***, p<0.001 versus untreated. (**C**) Human serous EC ARK1 and A375 melanoma cells were transfected with control nontargeting shRNA (ShNT) or ERK5-specific shRNAs (ShERK5-1 and ShERK5-2). After 5 days cells were lysed and Western Blot was performed with the indicated antibodies. Representative images from two independent experiments are shown. Molecular weight markers are indicated on the left.

### ERK5 inhibition induces CDK6 proteasomal degradation

Although CDK protein levels stay relatively constant during the cell cycle, they are continuously renewed through a balance between synthesis and ubiquitin-mediated proteasomal degradation^35,36^. Therefore, we asked whether the decrease in CDK6 observed upon ERK5 inhibition could result from increased proteasomal degradation. Notably, the proteasome inhibitor MG-132 prevented the degradation of CDK6 protein induced by the ERK5i JWG-071 in all cell lines tested (Fig. 2A-B). These results indicate that ERK5 kinase activity contributes to the stabilization of CDK6 protein, and that ERK5 inhibition promotes proteasome-dependent CDK6 degradation.

**Figure 2.**
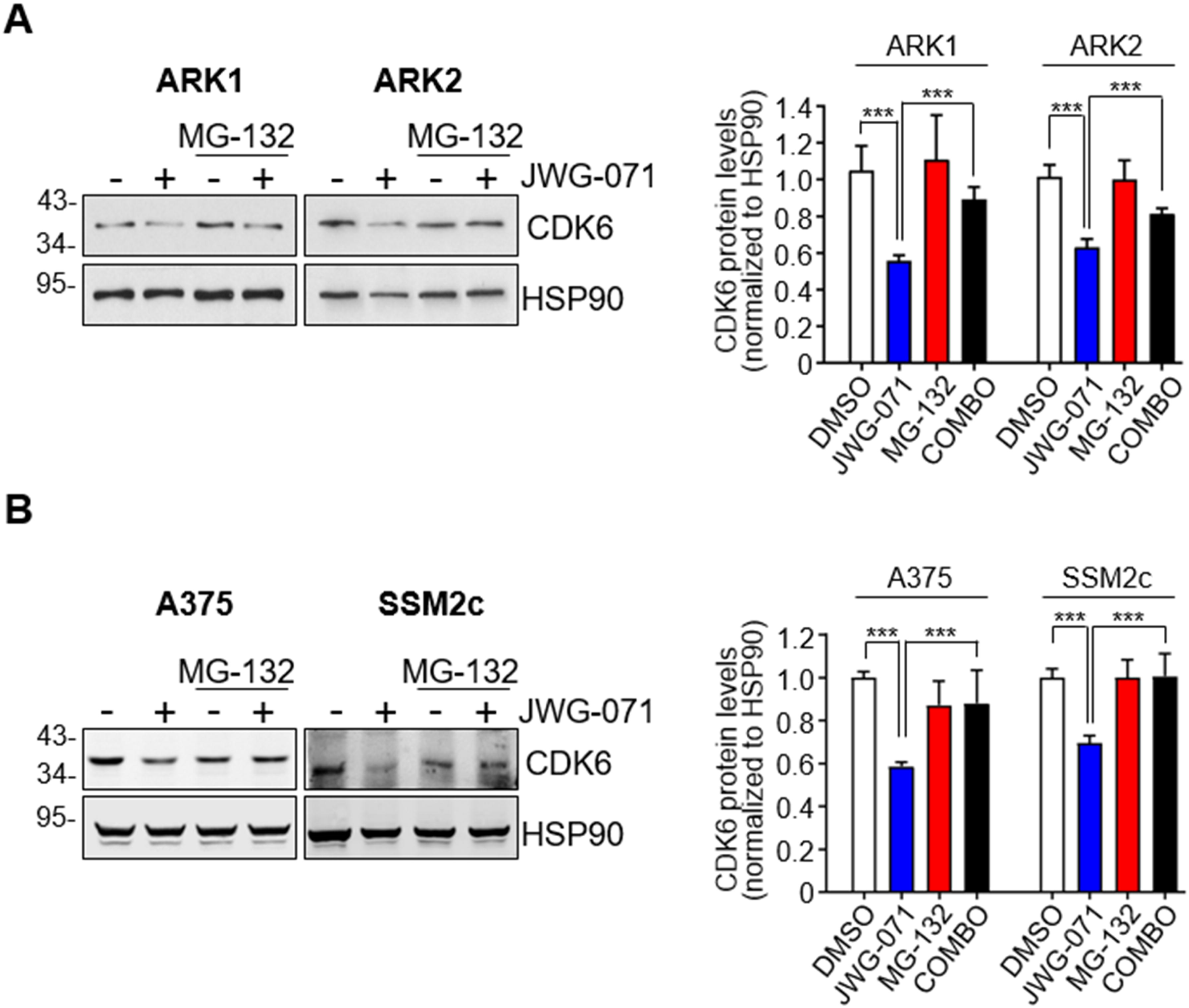
ERK5 inhibition induces CDK6 proteasomal degradation. (**A, B**) Human serous EC ARK1 and ARK2 cells (A) or human melanoma A375 and SSM2c cells (B) were treated with 5μM JWG-071 for 24 hours and with 10μM of the proteasome inhibitor MG-132 for the last 8 hours of the ERK5i treatment. Cells were then lysed and Western Blot was performed with the indicated antibodies. Molecular weight markers are indicated on the left. The graphs show average densitometric values normalized on HSP90 content ± SD of three independent experiments. One-way ANOVA variance analysis was used to assess statistical significance ***, p<0.001 versus untreated.

### ERK5 targeting synergizes with palbociclib to impair cell viability

Given that inhibition of ERK5 reduces CDK6 protein levels, we next explored whether targeting ERK5 could enhance the cytotoxic effects of the CDK4/6 inhibitor palbociclib used in the clinics^16^. To this end, MTT assays were performed using various combinations of an ERK5 inhibitor and palbociclib in human serous EC and melanoma cells. Low doses of JWG-071 significantly potentiated the effect of palbociclib in reducing cell viability, having little or no effect *per se* (Fig. 3A and B). BLISS synergy analysis rendered a strong synergistic interaction between the ERK5 inhibitor and palbociclib across several concentrations (Fig. 3C, BLISS score>10). Notably, this effect was consistent among cell lines with distinct molecular backgrounds, indicating that ERK5 inhibition potently synergizes with palbociclib to reduce cell viability independently of specific genetic contexts.

**Figure 3.**
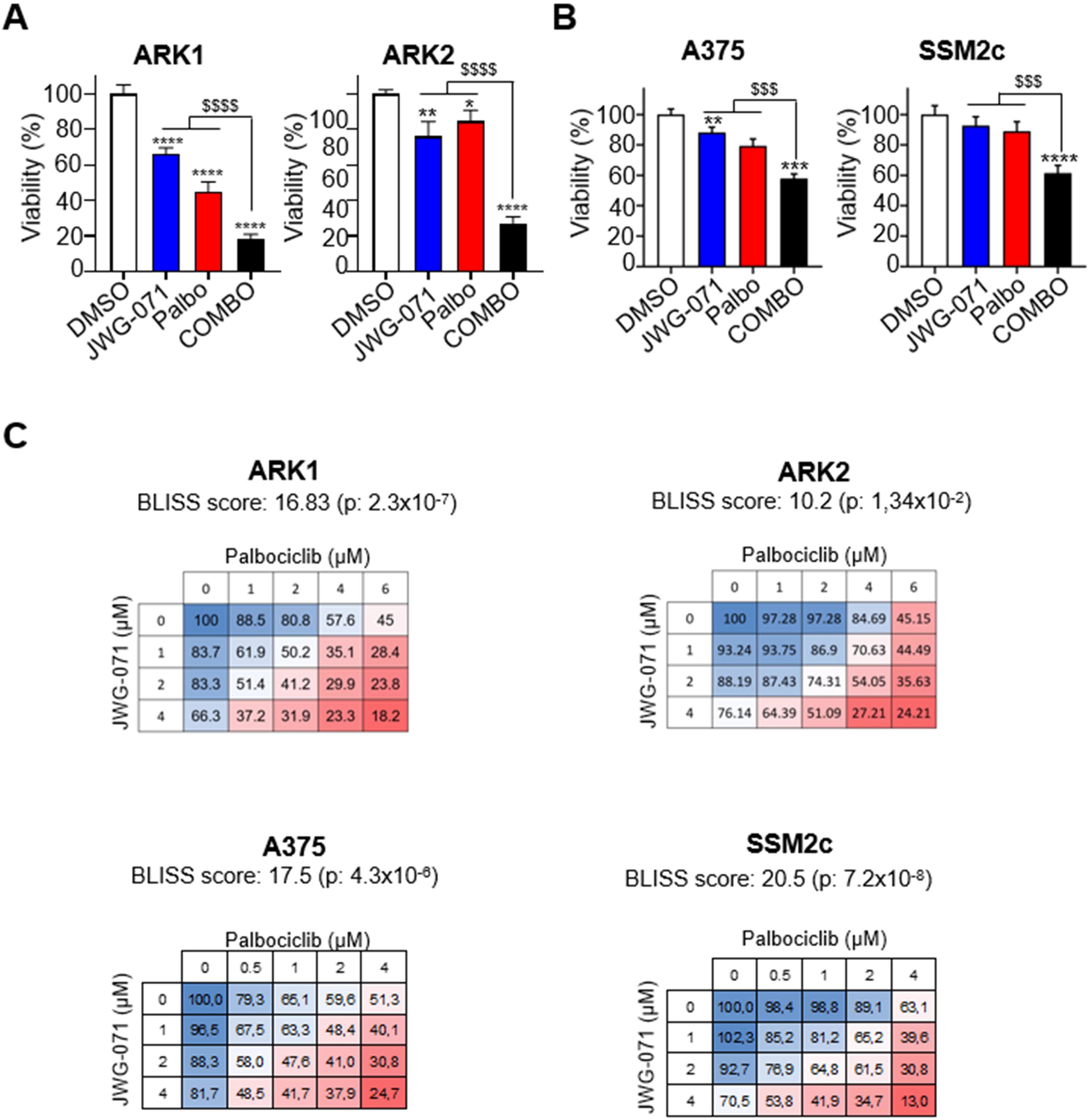
ERK5 inhibition synergizes with palbociclib in reducing cell viability. (**A-B**) Human serous EC ARK1 and ARK2 cells were treated with 5μM JWG-071 and/or 5μM palbociclib (A), human melanoma A375 were treated with 2 μM JWG-071 and/or 0.5μM palbociclib, and SSM2c cells were treated with 1μM JWG-071 and/or 1μM palbociclib (B). The effect on cell viability was evaluated with MTT assay after 72 hours. Graphs represent the mean percentage of viable cells ± SD of three independent experiments. One-way ANOVA variance analysis was used to assess statistical significance * p<0.05, ** p< 0.01, ***, p<0.001, ****, p<0.0001 versus untreated; ^$$$^ p< 0.001, ^$$$$^ p< 0.0001 among indicated conditions. (C) Cells were treated with different combinations of palbociclib and JWG-071 as indicated for 72 hours and MTT assay was then performed. Graphs represent the synergy score. A Bliss score higher than 10 is indicative of synergism.

### ERK5 inhibition potentiates the cytotoxicity of palbociclib

Since palbociclib primarily acts by blocking CDK4/6-dependent cell cycle progression at the G1/S transition, we next evaluated whether its combination with ERK5 inhibition translated into a reduced number of viable cells. Consistent with the findings of Figure 3, combined treatment of the ERK5 inhibitor JWG-071 and palbociclib resulted in a significantly lower number of EC and melanoma cells compared to either treatment alone (Fig. 4A and B), indicating that ERK5 inhibition potentiates the anti-proliferative activity of palbociclib across both tumor models.

**Figure 4.**
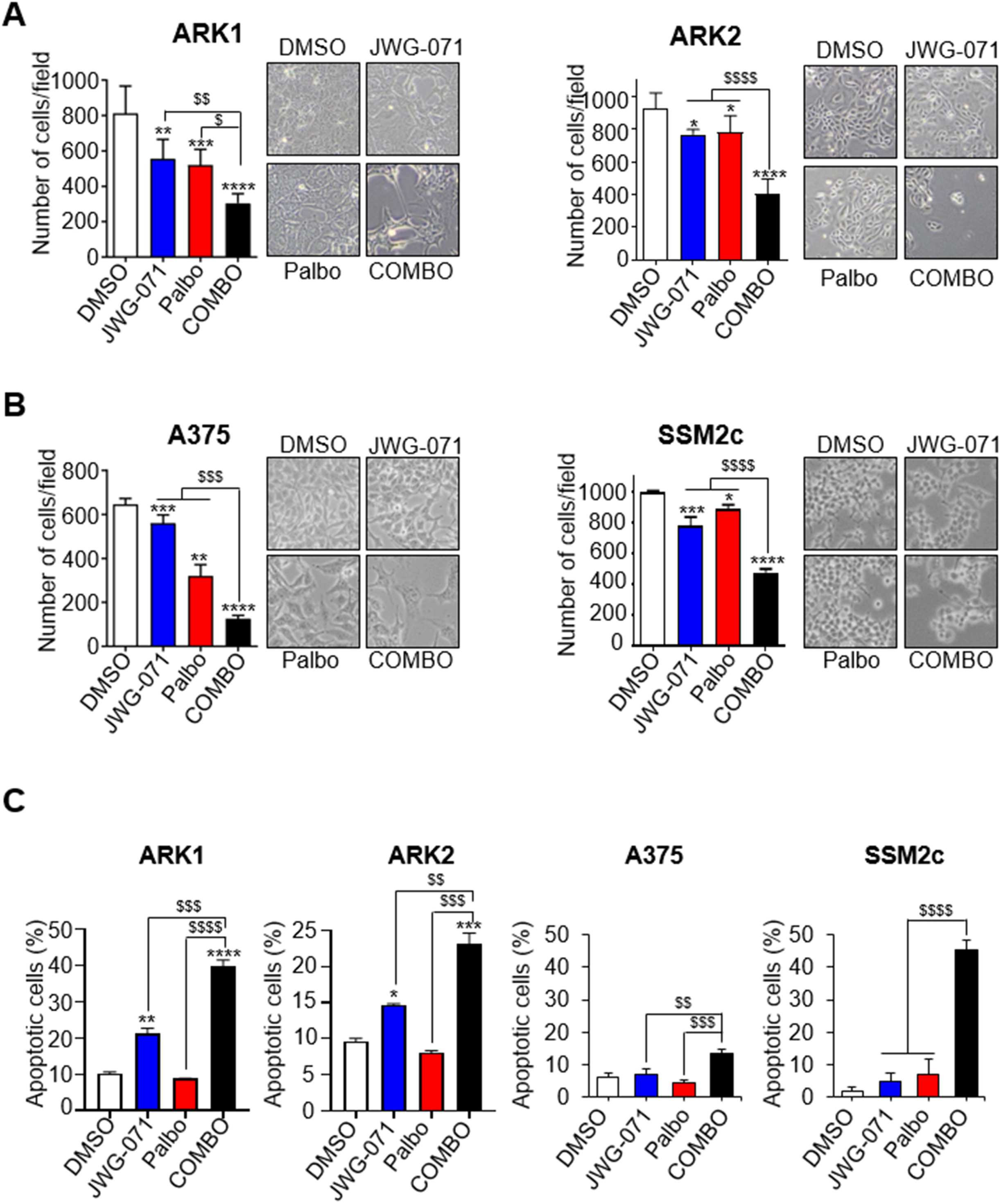
ERK5 inhibition potentiates the cytotoxicity of palbociclib. (**A, B**) Human serous EC were treated with vehicle (DMSO), 5μM JWG-071 and/or 5μM palbociclib for 24 hours (A), while A375 and SSM2c melanoma cells were treated with 1μM palbociclib and/or 1μM JWG-071 for 48 hours (B). Cells were visualized by phase-contrast microscopy in a cell counting assay. Histograms show the number of cells per field (3 fields per experimental condition) ± SD from three independent experiments. Representative images taken at 10x magnification are shown. One-way ANOVA variance analysis was used to assess statistical significance * p<0.05, ** p< 0.01, ***, p<0.001, ****, p<0.0001 versus untreated; ^$^ p< 0.05, ^$$^ p< 0.01, ^$$$^ p< 0.001, ^$$$$^ p< 0.0001 among indicated conditions. (**C**) ARK1 and ARK2 cells were treated with vehicle (DMSO), 5μM JWG-071 and/or 5μM palbociclib for 48 hours, while A375 and SSM2c cells were treated with vehicle (DMSO), 4μM JWG-071 and/or 2μM palbociclib for 48 hours. Apoptosis was evaluated through flow cytometry following Annexin V-propidium iodide staining. Graphs show the average percentage ± SD of apoptotic (annexin V-positive) cells in three independent experiments One-way ANOVA variance analysis was used to assess statistical significance. ** p< 0.01, ***, p<0.001 and ****, p<0.00001 versus untreated, ^$$^ p< 0.01, ^$$$^ p< 0.001, ^$$$$^ p< 0.0001 among indicated conditions.

To determine whether this reduced cell number was associated with increased cell death, we performed flow cytometric apoptosis analysis using Annexin V/propidium iodide staining. Co-treatment of cells with the ERK5 inhibitor and palbociclib significantly increased apoptosis in both EC and melanoma cells, compared with single-agent treatments (Fig. 4C), in agreement with the MTT and cell counting results.

To gain insight into the molecular basis of the cooperative effect impairing cell proliferation, we analyzed key cell cycle regulators involved in G1/S phase transition by immunoblot. Interestingly, the reduction of CDK6 protein levels observed upon ERK5 inhibition were maintained (A375) or further enhanced (ARK1) upon combined treatment with palbociclib (Figure 5 A and B). As control, palbociclib impaired the levels of phosphorylated Rb, a canonical substrate of CDK6, further indicating that the serous EC and melanoma cells used are sensitive to CDK4/6 inhibitors (Supplementary Figure 2). Interestingly, in both cellular models, the levels of phosphorylated Rb were further reduced upon co-treatment with JWG-071 and palbociclib. Of note, ERK5 inhibition resulted in increased p21 protein levels, as extensively reported^14,25^, and this effect was enhanced by the combined treatment in both cellular models, despite not reaching statistical significance in ARK1 cells (Fig. 5 A and B).

**Figure 5.**
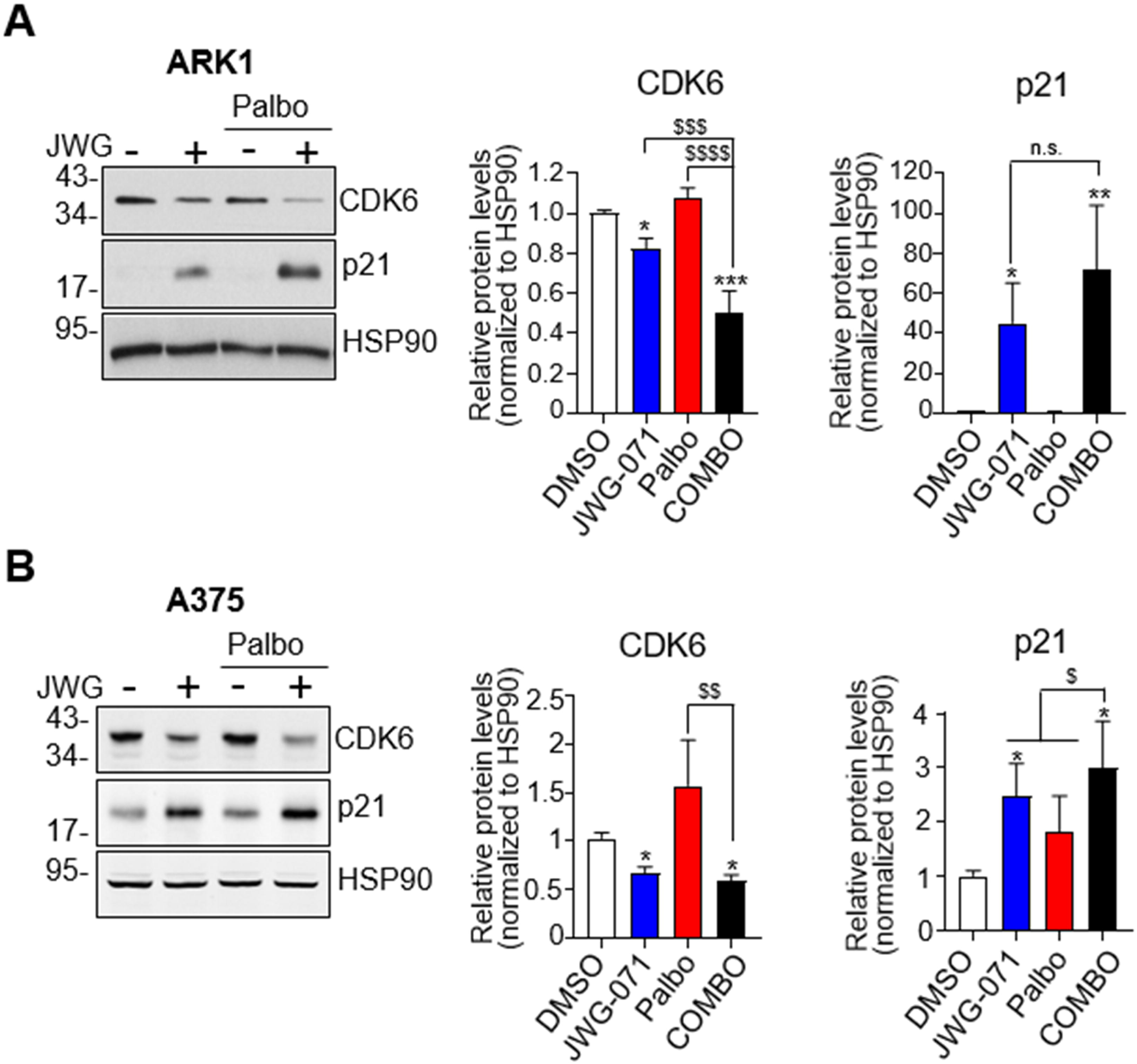
ERK5 inhibition combined with palbociclib treatment enhances p21 upregulation. (**A**) ARK1 cells were treated with vehicle (DMSO), 5μM JWG-071 and/or 5μM palbociclib for 48 hours, while A375 cells were treated with vehicle (DMSO), 4μM JWG-071 and/or 2μM palbociclib for 48 hours. The expression of the indicated proteins was analyzed by immunoblot. Molecular weight markers are indicated on the left. The graphs show average densitometric values normalized on HSP90 level ± SD of three independent experiments. One-way ANOVA variance analysis was used to assess statistical significance *, p<0.05, **, p<0.01, ***, p<0.001, ****, p<0.0001 versus untreated, ^$^ p< 0.05, ^$$^ p< 0.01, ^$$$$^ p< 0.0001 among indicated conditions. N.s. not significant.

Collectively, these results demonstrate that ERK5 inhibition synergizes with palbociclib to suppress cell proliferation and promote apoptosis in high-risk cancer cells, independently of their genetic background.

## DISCUSSION

Melanoma and serous EC are two malignancies with high mortality rates at advanced stages^37–40^. The CDK4/CDK6 inhibitor palbociclib is clinically approved and shows high efficacy in certain molecular cancer subtypes, and it is currently being investigated for other types of cancer, including EC and melanoma^3,4,41^. However, both adverse effects and the development of resistance have been reported^42–45^. In this regard, combination strategies with other small-molecule inhibitors that enhance the efficacy of palbociclib have been proposed in order to allow dose reduction and limit the occurrence of resistance^45,46^.

In this study, we evaluated the effects of combining palbociclib with a small molecule ERK5 inhibitor (namely JWG-071), and demonstrated that ERK5 inhibition potentiates the cytotoxic activity of palbociclib, supporting this combination as a promising therapeutic strategy for cancer treatment. Of relevance, we found that ERK5 inhibition impairs CDK6, but not CDK4, protein levels. The absence of effect of ERK5 inhibition on CDK4 is in line with a previous report performed using ERK5 knock-out MEF^26^. Our findings demonstrate that CDK6 is degraded by the proteasome upon ERK5 inhibition. Interestingly, ERK5 regulation of CDK6 proteostasis occurs irrespective of the tumor type and the molecular background. In fact, A375 cells carry BRAF V600E-mutation, SSM2c are triple wild-type (i.e. do not harbor BRAF, NRAS nor NF-1 mutations), whereas serous EC ARK1/2 cells are PI3KCA-mutated.

Several works have shown that ERK5 regulates protein stability through phosphorylation-dependent mechanisms. ERK5-mediated phosphorylation prevents the ubiquitylation and proteasomal degradation of c-Myc protein in pancreatic cancer cells ^47^, of KLF2 in mouse embryonic stem cells^48^, and of TP53INP2 in diverse cancer cell types^22^. In line with these observations, one plausible mechanism involved in CDK6 stabilization is that ERK5 stabilizes CDK6 by phosphorylation, so ERK5 inhibition may disrupt this stabilizing phosphorylation, thereby leading to proteasomal degradation of CDK6. This hypothesis is supported by the fact that ERK5 kinase inhibition using a number of ERK5i elicits the reduction of CDK6 protein levels.

A key finding of this study is that ERK5 inhibition enhances the cytotoxic effect of palbociclib in cancer cells. Notably, the ERK5 inhibitor JWG-071 and palbociclib exhibit a synergistic effect at low concentrations, leading to a significant reduction in cell viability in both melanoma and serous EC models. With regards to the molecular mechanisms underlying this synergism, we found that the combined treatment with JWG-071 and palbociclib further decreases or maintains CDK6 protein levels compared to ERK5 inhibition alone, depending on the cellular model, while eliciting an increase of p21, suggesting that the enhanced cell-cycle inhibitory signaling may contribute to the increased cytotoxic effect. Furthermore, combined ERK5-CDK4/6 inhibition resulted in a more pronounced reduction of the phosphorylation of Rb than the single treatments alone, highlighting a suppression of Rb-dependent cell-cycle progression. These findings suggest that ERK5 inhibition cooperates with palbociclib through many cell cycle regulators.

ERK5 inhibition has consistently been shown to potentiate the efficacy of chemotherapy across multiple solid tumor types^12,21,49–51^. Our findings extend this paradigm by identifying ERK5 targeting as a rational second-hit strategy to enhance tumor vulnerability by potentiating the activity of CDK4/6 blockade through palbociclib. These data provide mechanistic and translational support for ERK5 inhibition as a combinatorial approach to reinforce cell cycle-directed therapies. Future investigations, incorporating additional clinically approved CDK4/6 inhibitors, such as abemaciclib and ribociclib, will be instrumental in defining the broader therapeutic scope of ERK5-based combinations.

## Author contributions

S. Espinosa-Gil and E. Rovida conceived and designed the research; S. Espinosa-Gil, Y. Sfragano, C. Faundez-Vidiella and I. Tusa performed the research and acquired the data; B. Stecca provided reagents and competences; Y. Sfragano, C. Faundez-Vidiella, I. Tusa, S. Espinosa-Gil, J. Lizcano, and E. Rovida analyzed and interpreted the data. Y. Sfragano and C. Faundez-Vidiella wrote the draft of the manuscript; E. Rovida and J. Lizcano edited and revised the manuscript.

## Data availability statement

All data generated or analyzed during this study are included in this published article and are available from the corresponding authors upon reasonable request.

## Acknowledgments

The experimental procedures performed in ER’s lab took advantage of the Molecular Medicine Facility of the Department of Experimental and Clinical Biomedical Sciences, supported in part by the Italian Ministry of Education, University and Research (MIUR). The research leading to these results has received funding from AIRC under IG 2025-ID. 32014 project–P.I. Rovida Elisabetta, European Union NextGenerationEU-National Recovery and Resilience Plan (NRRP) ref. ECS00000017-CUP B83C22003920001, and from Fondo Beneficenza Intesa San Paolo. JML laboratory is supported by the Spanish Ministry of Science and Innovation (grant references PID2019-107561RB-I00 and PID2022-136391OB-I00), and co-funded by the European Regional Development Fund (ERDF, “A way to make Europe/Investing in your future”). CF-V is supported by a grant from the Scientific Foundation of the Spanish Association Against Cancer (Ref. PRDBA246090FAUN). YS is supported by NRRP (D.M. 118/2023)/NEXTGENERATIONEU. SE-G was a recipient of fellowships from FI-AGAUR of the Generalitat de Catalunya (2020-FISDU-00575 and FI-B-00293).

---

**Supplementary Figure 1. Treatment with ERK5 inhibitors induce a reduction in CDK6 protein levels.** (**A**, **B**) Human serous EC ARK1 and ARK2 (**A**) or human melanoma A375 (**B**) cells were treated with of the indicated drugs for 24 hours. Cells were then lysed and Western Blot was performed with the indicated antibodies. Molecular weight markers are indicated on the left.

**Supplementary Figure 2. Effect of combined JWG-071 and palbociclib on RB phosphorylation.** (**A**, **B**) ARK1 cells were treated with vehicle (DMSO), 5μM JWG-071 and/or 5μM palbociclib for 48 hours (A), while A375 cells were treated with vehicle (DMSO), 5 μM JWG-071 and/or 10μM palbociclib for 48 hours. The expression of the indicated proteins was analyzed by immunoblot and reported in the graphs as average densitometric values normalized on Hsp90 content ± SD of three (**A**) or two (**B**) independent experiments. One-way ANOVA variance analysis (**A**) or t test (**B**) were used to assess statistical significance *, p<0.05, **, p<0.01, ***, p<0.001 versus untreated, $ p< 0.05, $$ p< 0.01 among indicated conditions.

## REFERENCES

1. Henley, S. J. et al. Annual report to the nation on the status of cancer, part I: National cancer statistics. Cancer 126, 2225–2249 (2020).

2. Sung, H. et al. Global Cancer Statistics 2020: GLOBOCAN Estimates of Incidence and Mortality Worldwide for 36 Cancers in 185 Countries. CA Cancer J. Clin. 71, 209–249 (2021).

3. Comstock, C. E. S. et al. Targeting cell cycle and hormone receptor pathways in cancer. Oncogene 32, 5481–5491 (2013).

4. Powell, M. A. et al. Efficacy and safety of dostarlimab in combination with chemotherapy in patients with dMMR/MSI-H primary advanced or recurrent endometrial cancer in a phase 3, randomized, placebo-controlled trial (ENGOT-EN6-NSGO/GOG-3031/RUBY). Gynecol. Oncol. 192, 40–49 (2025).

5. Brinton, L. A. et al. Etiologic heterogeneity in endometrial cancer: Evidence from a Gynecologic Oncology Group trial. Gynecol. Oncol. 129, 277–284 (2013).

6. Davis, L. E., Shalin, S. C. & Tackett, A. J. Current state of melanoma diagnosis and treatment. Cancer Biol. Ther. 20, 1366–1379 (2019).

7. Guo, L., Qi, J., Wang, H., Jiang, X. & Liu, Y. Getting under the skin: The role of CDK4/6 in melanomas. Eur. J. Med. Chem. 204, 112531 (2020).

8. Kim, U. et al. Advances in Cutaneous Melanoma Therapy: The Emerging Role of CDK4/6 Inhibitors. Pharmacol. Res. 221, 107955 (2025).

9. Kato, Y. et al. Bmk1/Erk5 is required for cell proliferation induced by epidermal growth factor. Nature 395, 713–716 (1998).

10. Stecca, B. & Rovida, E. Impact of ERK5 on the Hallmarks of Cancer. Int. J. Mol. Sci. 20, 1426 (2019).

11. McCracken, S. R. C. et al. Aberrant expression of extracellular signal-regulated kinase 5 in human prostate cancer. Oncogene 27, 2978–2988 (2008).

12. Diéguez-Martínez, N. et al. The ERK5/NF-κB signaling pathway targets endometrial cancer proliferation and survival. Cellular and Molecular Life Sciences 79, 524 (2022).

13. Montero, J. C. et al. Expression of Erk5 in Early Stage Breast Cancer and Association with Disease Free Survival Identifies this Kinase as a Potential Therapeutic Target. PLoS One 4, e5565 (2009).

14. Tusa, I. et al. ERK5 is activated by oncogenic BRAF and promotes melanoma growth. Oncogene 37, 2601–2614 (2018).

15. Sánchez-Fdez, A. et al. MEK5 promotes lung adenocarcinoma. European Respiratory Journal 53, 1801327 (2019).

16. Espinosa-Gil, S. et al. MAP kinase ERK5 modulates cancer cell sensitivity to extrinsic apoptosis induced by death-receptor agonists and Natural Killer cells. Preprint at 10.1101/2023.03.22.533738 (2023).

17. Pereira, D. M. & Rodrigues, C. M. P. Targeted Avenues for Cancer Treatment: The MEK5–ERK5 Signaling Pathway. Trends Mol. Med. 26, 394–407 (2020).

18. Tubita, A., Tusa, I. & Rovida, E. Playing the Whack-A-Mole Game: ERK5 Activation Emerges Among the Resistance Mechanisms to RAF-MEK1/2-ERK1/2-Targeted Therapy. Front. Cell Dev. Biol. 9, (2021).

19. Tusa, I., et al. The Hedgehog-GLI Pathway Regulates MEK5-ERK5 Expression and Activation in Melanoma Cells. Int. J. Mol. Sci. 22, 11259 (2021).

20. Tusa, I. et al. The MEK5/ERK5 pathway promotes the activation of the Hedgehog/GLI signaling in melanoma cells. Cellular Oncology 48, 789–799 (2025).

21. Pereira, D. M. et al. MEK5/ERK5 signaling inhibition increases colon cancer cell sensitivity to 5-fluorouracil through a p53-dependent mechanism. Oncotarget 7, 34322–34340 (2016).

22. Espinosa-Gil, S. et al. MAP kinase ERK5 modulates cancer cell sensitivity to extrinsic apoptosis induced by death-receptor agonists. Cell Death Dis. 14, 715 (2023).

23. Mulloy, R., Salinas, S., Philips, A. & Hipskind, R. A. Activation of cyclin D1 expression by the ERK5 cascade. Oncogene 22, 5387–5398 (2003).

24. Rovida, E. et al. The mitogen-activated protein kinase ERK5 regulates the development and growth of hepatocellular carcinoma. Gut 64, 1454–1465 (2015).

25. Rovida, E. et al. ERK5/BMK1 Is Indispensable for Optimal Colony-Stimulating Factor 1 (CSF-1)-Induced Proliferation in Macrophages in a Src-Dependent Fashion. The Journal of Immunology 180, 4166–4172 (2008).

26. Perez-Madrigal, D., Finegan, K. G., Paramo, B. & Tournier, C. The extracellular-regulated protein kinase 5 (ERK5) promotes cell proliferation through the down-regulation of inhibitors of cyclin dependent protein kinases (CDKs). Cell. Signal. 24, 2360–2368 (2012).

27. Tubita, A. et al. Inhibition of ERK5 Elicits Cellular Senescence in Melanoma via the Cyclin-Dependent Kinase Inhibitor p21. Cancer Res. 82, 447–457 (2022).

28. Paudel, R. et al. MEK5/ERK5 inhibition sensitizes NRAS-mutant melanoma to MAPK-targeted therapy by preventing Cyclin D/CDK4-mediated G1/S progression. Cell Death Dis. 16, 689 (2025).

29. Pandolfi, S. et al. WIP1 phosphatase modulates the Hedgehog signaling by enhancing GLI1 function. Oncogene 32, 4737–4747 (2013).

30. Wang, J. et al. Structural and Atropisomeric Factors Governing the Selectivity of Pyrimido-benzodiazipinones as Inhibitors of Kinases and Bromodomains. ACS Chem. Biol. 13, 2438–2448 (2018).

31. Bono, S. et al. Different BCR/Abl protein suppression patterns as a converging trait of chronic myeloid leukemia cell adaptation to energy restriction. Oncotarget 7, 84810–84825 (2016).

32. Gámez-García, A. et al. ERK5 Inhibition Induces Autophagy-Mediated Cancer Cell Death by Activating ER Stress. Front. Cell Dev. Biol. 9, 742049 (2021).

33. Cheloni, G. et al. The Leukemic Stem Cell Niche: Adaptation to “Hypoxia” versus Oncogene Addiction. Stem Cells Int. 2017, 1–8 (2017).

34. Barbetti, V., Tusa, I., Cipolleschi, M. G., Rovida, E. & Dello Sbarba, P. AML1/ETO sensitizes via TRAIL acute myeloid leukemia cells to the pro-apoptotic effects of hypoxia. Cell Death Dis. 4, e536–e536 (2013).

35. Wang, H. C/EBPalpha triggers proteasome-dependent degradation of cdk4 during growth arrest. EMBO J. 21, 930–941 (2002).

36. Iizumi, Y. et al. Stabilization of CDK6 by ribosomal protein uS7, a target protein of the natural product fucoxanthinol. *Commun*. Biol. 5, 564 (2022).

37. Davis, L. E., Shalin, S. C. & Tackett, A. J. Current state of melanoma diagnosis and treatment. Cancer Biol. Ther. 20, 1366–1379 (2019).

38. Cronin, K. A. et al. Annual report to the nation on the status of cancer, part 1: National cancer statistics. Cancer 128, 4251–4284 (2022).

39. Murali, R., Soslow, R. A. & Weigelt, B. Classification of endometrial carcinoma: more than two types. Lancet Oncol. 15, e268–e278 (2014).

40. Bray F et al. Erratum: Global cancer statistics 2018: GLOBOCAN estimates of incidence and mortality worldwide for 36 cancers in 185 countries. CA Cancer J. Clin. 70, 313–313 (2020).

41. Mao, L. et al. Palbociclib in advanced acral melanoma with genetic aberrations in the cyclin-dependent kinase 4 pathway. Eur. J. Cancer 148, 297–306 (2021).

42. Zhang, S., Zhang, R., Xia, Y. & Cao, D. Safety assessment of cyclin-dependent kinase 4/6 inhibitors and comparison of time to adverse events. PLoS One 20, e0336767 (2025).

43. Kiener, T., Li, K. H., Chiang, E. & Le, Q. A. Impact of CDK4/6 inhibitors on health-related quality of life outcomes in patients with metastatic breast cancer: A systematic review and meta-analysis. J. Manag. Care Spec. Pharm. 31, 1285–1303 (2025).

44. Li, T. et al. Cyclin-dependent kinase 4/6 inhibitors and cardiotoxic events in breast cancer: A pharmacovigilance study based on the FAERS database. Int. J. Cancer 156, 1404–1418 (2025).

45. Scheidemann, E. R. & Shajahan-Haq, A. N. Resistance to CDK4/6 Inhibitors in Estrogen Receptor-Positive Breast Cancer. Int. J. Mol. Sci. 22, 12292 (2021).

46. Choi, P. J. et al. Conjugation of Palbociclib with MHI-148 Has an Increased Cytotoxic Effect for Breast Cancer Cells and an Altered Mechanism of Action. Molecules 27, 880 (2022).

47. Vaseva, A. V. et al. KRAS Suppression-Induced Degradation of MYC Is Antagonized by a MEK5-ERK5 Compensatory Mechanism. Cancer Cell 34, 807–822.e7 (2018).

48. Brown, H. A. et al. An ERK5–KLF2 signalling module regulates early embryonic gene expression and telomere rejuvenation in stem cells. Biochemical Journal 478, 4119–4136 (2021).

49. Song, C., et al. Correction: Corrigendum: Targeting BMK1 Impairs the Drug Resistance to Combined Inhibition of BRAF and MEK1/2 in Melanoma. Sci. Rep. 7, 46833 (2017).

50. Adam, C. et al. Efficient Suppression of NRAS-Driven Melanoma by Co-Inhibition of ERK1/2 and ERK5 MAPK Pathways. Journal of Investigative Dermatology 140, 2455–2465.e10 (2020).

51. Carmell, N. et al. Identification and Validation of ERK5 as a DNA Damage Modulating Drug Target in Glioblastoma. Cancers (Basel). 13, 944 (2021).

