## Supplementary material for "ERK5 inhibition triggers CDK6 proteasomal degradation and enhances palbociclib efficacy in cancer cells": Table 1

**Table 1**. List of antibodies used for Western Blot.

| CDK4 | Rabbit monoclonal | #12790 | Cell Signaling Technology, Danvers, MA, USA | RRID: AB_2631166 |
| --- | --- | --- | --- | --- |
| CDK6 | Mouse monoclonal | #3136 | Cell Signaling Technology, Danvers, MA, USA | RRID: AB_2229289 |
| ERK5 | Rabbit polyclonal | #3372 | Cell Signaling Technology, Danvers, MA, USA | RRID: AB_330491 |
| Hsp90 | Mouse monoclonal | sc-13119 | Santa Cruz Biotechnology, Santa Cruz, CA, USA | RRID: AB_675659 |
| p21Waf1/Cip1 | Rabbit monoclonal | #2947 | Cell Signaling Technology, Danvers, MA, USA | RRID: AB_823586 |
| pRb | Rabbit monoclonal | #8516 | Cell Signaling Technology, Danvers, MA, USA | RRID: AB_11178658 |
