## Supplementary figures and images for "ERK5 inhibition triggers CDK6 proteasomal degradation and enhances palbociclib efficacy in cancer cells"

### Supplemental Figure 1

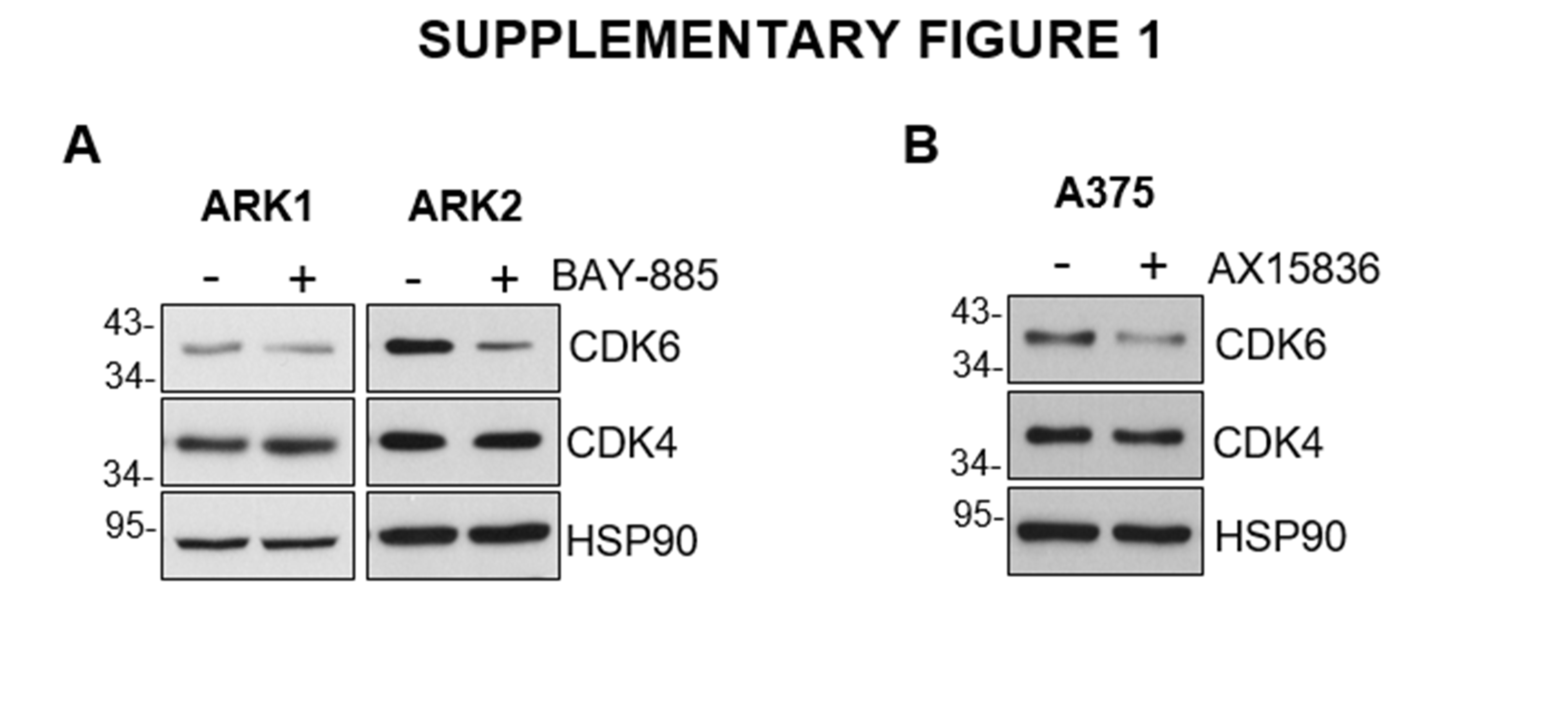

### Supplemental Figure 2

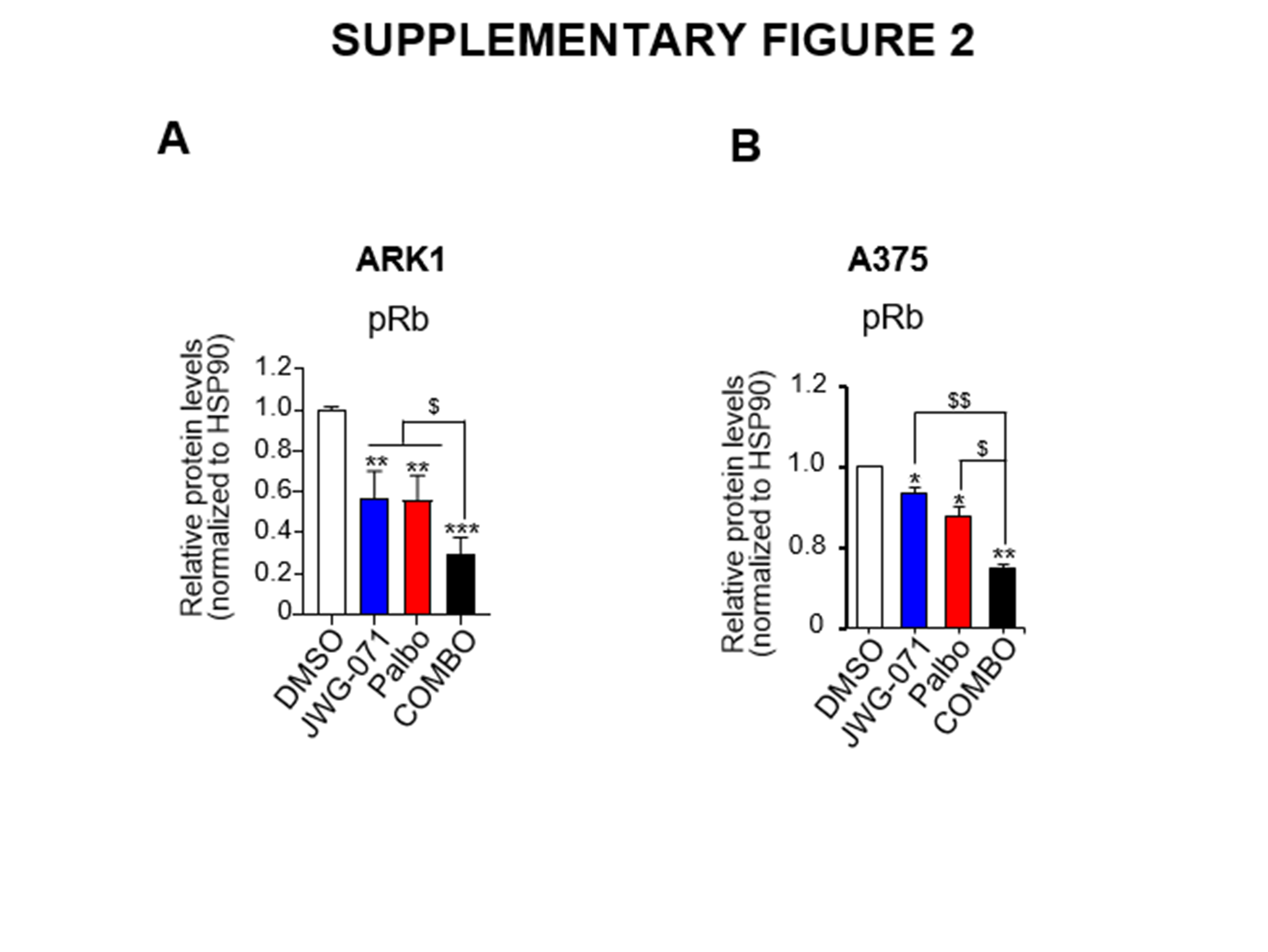
